# Seasonal diversity of soil microarthropods in two different vegetable plots of Aligarh-India

**DOI:** 10.1101/687483

**Authors:** Mohammad Jalaluddin Abbas, Hina Parwez

## Abstract

Soil microarthropods are intimately linked with health and fertility of soil as well as plant productivity. In India, despite their rich faunal diversity, information on soil microarthropods diversity and interactions with variety of edaphic factors is extremely limited. The present study has been carried out to observe seasonal diversity of soil microarthropods in two different vegetable plots at Aligarh. The two vegetable plots chosen in this study where predominantly Tomato (*Lycopersicom esculentum*) and Brinjal (*Solanum melongena*, family-Solaneceae) have been grown and sapling has been sown in the month of April when plants attained a height of approximately 6″. The samples were collected randomly from a depth of 5cm. @ of four samples per month for a period of one year. All microarthropods extracted with the help of Tullgren funnel apparatus. Among soil microarthropods collected, Collembolans have highest average monthly density (15.20 inds./sample) in brinjal plot and greatest abundance (18.7inds./sample) in tomato plot. A highly significant negative correlation was recorded between Collembolans population with reference to soil temperature (r = −0.867, P<0.05), whereas available nitrogen showed a positive correlation (r = 0.847, P>0.05). Interestingly, at neutral pH level, the highest population of Collembolans as well as Acari(mites) were recorded. During spring and winter months, there was a peak population buildup of Collembola and Acarina, whereas a sharp decline was recorded in summer months. So, this study clearly establishes that habitat difference as well as edaphic factors plays an important role along with seasonal parameters on their diversity.

## Introduction

Soil biodiversity can be evaluated by means of maximizing the numbers of reliable soil indicators that they exist in the soil as functional attributes in an ecosystem. Apart from the functional attributes of soil ecosystem, soil microarthropods make up a large and functionally important part of the soil biota (Siepel and Maaskamp 1994, Maraun et al. 2003) and they contribute significantly in decomposition by shredding and feeding on soil organic matter/fungi, and to release nutrients available (Teuben and Verhoef 1992, Siepel and Maaskamp 1994, Mc Gonigle 1995, Krivtsov et al. 2004, and Schneider et al. 2004) for plants. Thus, they help in promoting the plant growth either directly or indirectly. Among the soil microarthropods, Collembolans are vital source to sustain the soil health and they never cause diseases except very few cases.

Consequently, soil microarthropods such as Collembolans are good bio-indicators of arable soil quality (Rusek-1998), are very sensitive to variations in soil environment (Parisi-2001) and conditioning to detritus for microbial breakdown as well as farming soil to form soil micro-structure (Chahartaghi-2005). However, Collembola communities are still poorly understood (Alvarez et al. 2000) in spite of the role of these micro organisms in soil mineralization and humification of organic matter (Coleman 1985, Huhta et al. 1988, Czarnecki 1989, Striganova 1992) and they are considered as the bio-indicators in studies of soil quality (Heisler 1995, Kopeszki 1997) and soil health status.

In our study, results are highlighted to Collembolans and acarina (mites) and other soil dwelling microarthropods (Pterygotes) are minimized because both acari and Collembolans accounted approximately 75% of the total population of soil microarthropods. Thus, for the purpose of making contribution to the site variation, we carried out an experiment at Aligarh and compare the seasonal diversity as well as overall population dynamics of soil microarthropods with in two vegetational plots.

## Materials and Methods

### (a) Study Site

The site of investigation was situated at Zakir Bagh, which is at the heart of Aligarh Muslim University (AMU) campus. The boundary of site was shaded with higher plants such as Mango (*Mangifera indica*) and Neem (*Azadirachta indica*); however, these plants were 10 feet far from each of the plot boundary. Thus, both plots were semi shaded areas. For the purpose of soil samples, we considered a vegetation garden having two different plots of 5×5 meters of its area. Tomato (*Lycopersicom esculentum*) and Brinjal (*Solanum melongena*) vegetables (family-Solaneceae) were grown in the very beginning of March. The sapling were sown in the month of April when the plants attained a height of approximately 6″, soil samples were collected. Each plot has 25 plants of its vegetation. These vegetables have been selected, because they may grow throughout the year. The plants of Solaneceae family generally have more root-root network than compare to others. Thus, they play a significant role in soil aggregation and in maintaining the soil profile for better fertility and provide help in humus formation due to its plant litter material too. These plants grow between pH 6.5 to 7.5 that is the normal range of the soil of this region. The soil of the plots examined was alluvial type (a mixture of sand, silt and clay) and more fertile in terms of productive manner.

### (b) Sampling, extraction and identification of soil microarthropods

As mentioned earlier, samples were taken every week regularly and the points selected within the plots were distributed randomly. Total 96 samples have been taken for site study during the investigation period from both plots. Each sample consists of 4 corers of 5 cm. size. Modified Tullgren funnel apparatus was used for extraction of soil microarthropods. The power of bulbs used was 60 watts. All microarthropods were collected inside a beaker which contained 70 % alcohol with few drops of glycerol and they were separated and mounted with DPX. All soil microarthropods were identified up to the level of their order or, family using a range of taxonomic keys (O’ Connell and Bolger 1997). A binocular stereomicroscope was used to identify the soil microarthropods.

### (c) Statistical Analysis

To study the population dynamics of soil microarthropods, the parameters considered were density and abundance as stated previously (Abbas and Parwez, 2009) and fractional population dynamics, relative density, absolute frequency etc. One way analysis of variance (ANOVA) was calculated for the significance of population of soil microarthropods with edaphic factors at 5 % level of significance.

## Results

<u>In brinjal plot</u>, total 58.49% Apterygotes, 27.79% Pterygotes and 12.93% Acarina(mites) were recorded during the study period (table, 2). Among the soil dwelling Pterygotes, Hymenoptera was the highly dance taxa comprising by up to 60.0 % of total population followed by Dipterans (21.7%) and the rest of all taxa. The peak population of soil dwelling Pterygotes found in summer while sharp decline in winter months. Among the Apterygote soil microarthropods, Collembolans have the highest individual population (71.43 %) recorded in spring, while the lowest (5.90 %) population recorded in rainfall. The highest density and abundance of Collembolans found in March (60.50) respectively while the sharp declines in May & September (Table, 1). The peak population of Acari (57.66 %) recorded in spring while the least (11.65 %) recorded in summer months. Most of Acari were of astigmatid mites while some of them were mesostigmatids. Apart from the Apterygote soil microarthropods, Collembolans have the highest population (98.49 %) as recorded in our experiment (table, 2).

**Table: 1.**
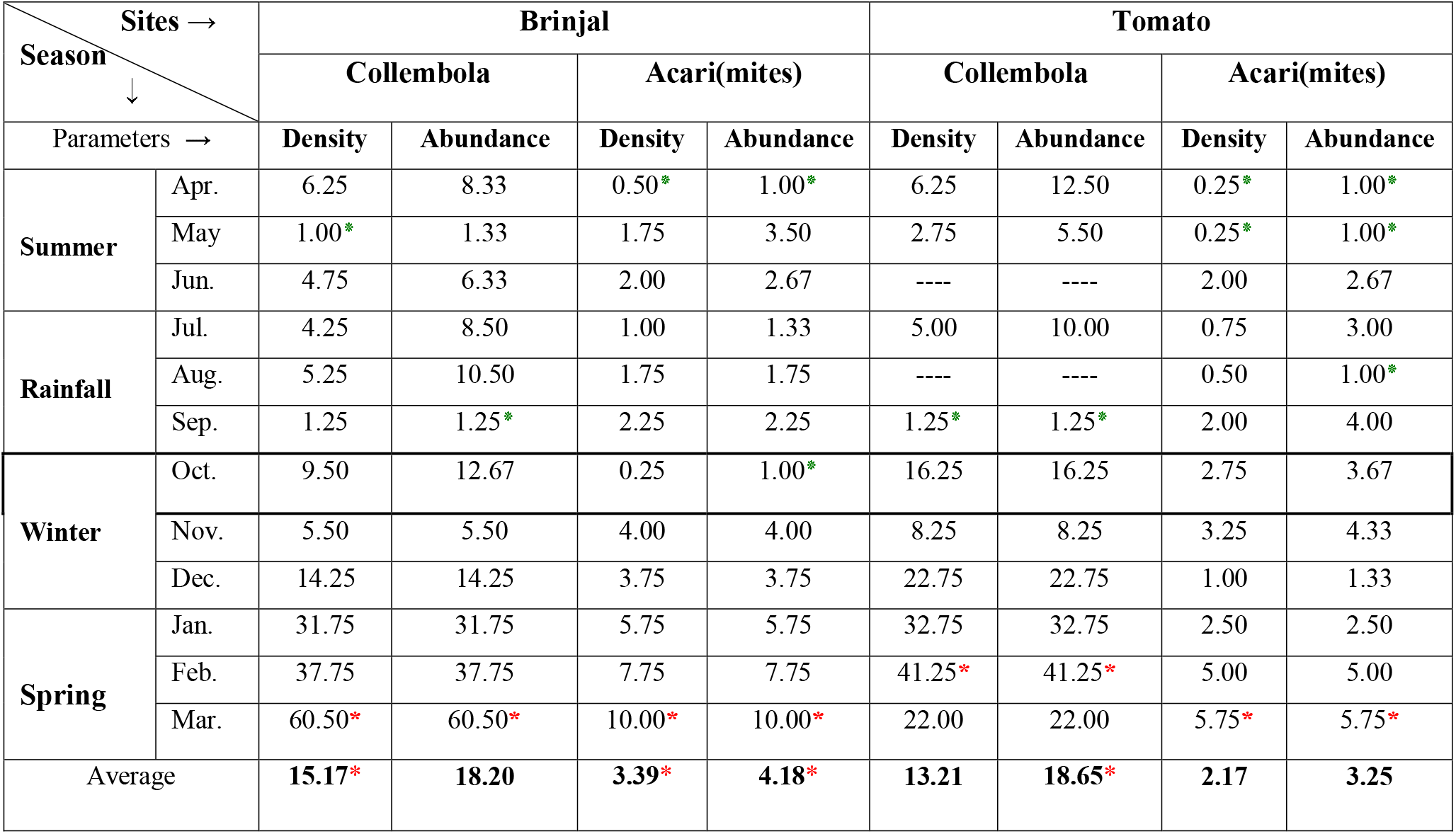
Seasonal variation of Collembola and Acari(mites) between two vegetational plots

| Sites →<br>Season ↓ |  | Brinjal |  |  |  | Tomato |  |  |  |
| --- | --- | --- | --- | --- | --- | --- | --- | --- | --- |
|  |  | Collembola |  | Acari(mites) |  | Collembola |  | Acari(mites) |  |
| Parameters → |  | Density | Abundance | Density | Abundance | Density | Abundance | Density | Abundance |
| Summer | Apr. | 6.25 | 8.33 | 0.50* | 1.00* | 6.25 | 12.50 | 0.25* | 1.00* |
|  | May | 1.00* | 1.33 | 1.75 | 3.50 | 2.75 | 5.50 | 0.25* | 1.00* |
|  | Jun. | 4.75 | 6.33 | 2.00 | 2.67 | ---- | ---- | 2.00 | 2.67 |
| Rainfall | Jul. | 4.25 | 8.50 | 1.00 | 1.33 | 5.00 | 10.00 | 0.75 | 3.00 |
|  | Aug. | 5.25 | 10.50 | 1.75 | 1.75 | ---- | ---- | 0.50 | 1.00* |
|  | Sep. | 1.25 | 1.25* | 2.25 | 2.25 | 1.25* | 1.25* | 2.00 | 4.00 |
| Winter | Oct. | 9.50 | 12.67 | 0.25 | 1.00* | 16.25 | 16.25 | 2.75 | 3.67 |
|  | Nov. | 5.50 | 5.50 | 4.00 | 4.00 | 8.25 | 8.25 | 3.25 | 4.33 |
|  | Dec. | 14.25 | 14.25 | 3.75 | 3.75 | 22.75 | 22.75 | 1.00 | 1.33 |
| Spring | Jan. | 31.75 | 31.75 | 5.75 | 5.75 | 32.75 | 32.75 | 2.50 | 2.50 |
|  | Feb. | 37.75 | 37.75 | 7.75 | 7.75 | 41.25* | 41.25* | 5.00 | 5.00 |
|  | Mar. | 60.50* | 60.50* | 10.00* | 10.00* | 22.00 | 22.00 | 5.75* | 5.75* |
| Average |  | 15.17* | 18.20 | 3.39* | 4.18* | 13.21 | 18.65* | 2.17 | 3.25 |

**Table: 2.** Relative density and Absolute frequency of total soil microarthropods within a year from two vegetational plots at Aligarh.

| Parameter → | Relative density (%) |  | Absolute frequency (%) |  |
| --- | --- | --- | --- | --- |
| Study sites → | Brinjal | Tomato | Brinjal | Tomato |
| Collembola | 57.77 | 52.83 | 83.33 | 70.83 |
| Protura | 1.43 | 1.43 | 12.50 | 12.50 |
| Diplura | 0.08 | ----- | 2.08 | ----- |
| Acari(mites) | 12.93 | 9.87 | 81.25 | 66.67 |
| Pterygotes | 27.79 | 35.87 | 77.08 | 87.50 |

<u>In tomato plot</u>, total 54.26 % Apterygotes, 35.87 % Pterygotes and 9.87 % Acari(mites) found with in a year (table, 2). Among the Aterygote soil microarthropods, Collembola was a highly diverse group (98.57 %) in terms of individual population (table 2). Among the soil dwelling Pterygotes, Hymenopterans were present about 43.63 % of entire Pterygotes community. The highest density and abundance of Collembolans found in February (41.25) respectively while the sharp declines in June & August (Table, 1).

There was significant difference in diversity of soil microarthropods between the plots investigated whereas the plots situated at one sequence. Brinjal plot was more diversified than the tomato plot. The average monthly abundance of Collembola was higher in tomato plot (18.65) than in brinjal plot (18.20); however, average density was lower in tomato plot (13.21) than in brinjal plot (15.17) as recorded during study period (Table, 1). Collembolans and Acari(mites) both were higher in most of the cases in brinjal plot than tomato plot (figure 1). The similar trends of peak population buildup of soil microarthropods in winter and spring months while sharp decline in the summer months in both kitchen plots have been recorded (Table 1). However, values of their diversity are quite different. Among soil microarthropods investigated, Collembolans were always found higher than Acari in both vegetable plots (figure, 1).

**Figure: 1.**
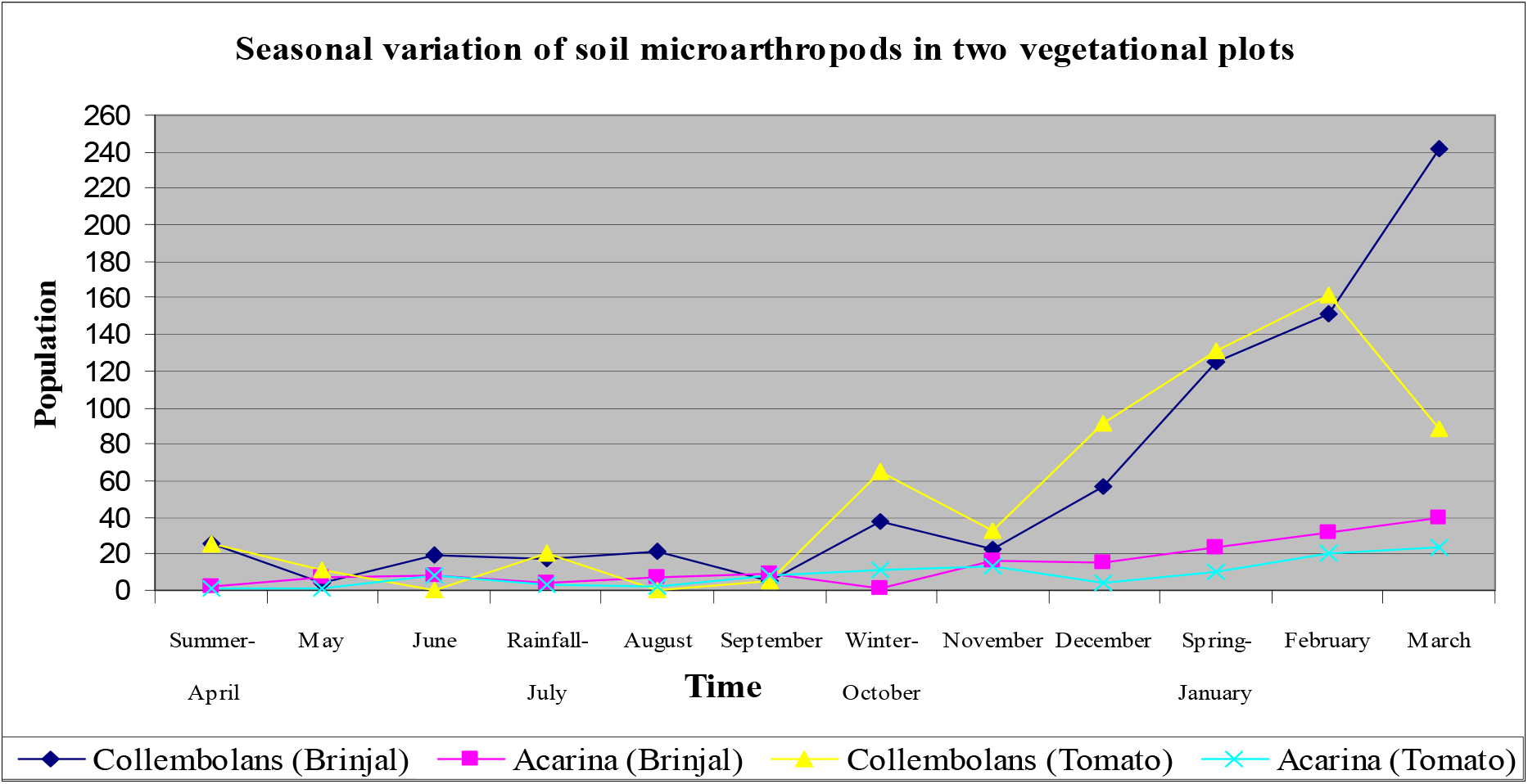
Seasonal diversity of soil microarthropods within two vegetational plots at Aligarh.

**Table: 3.**
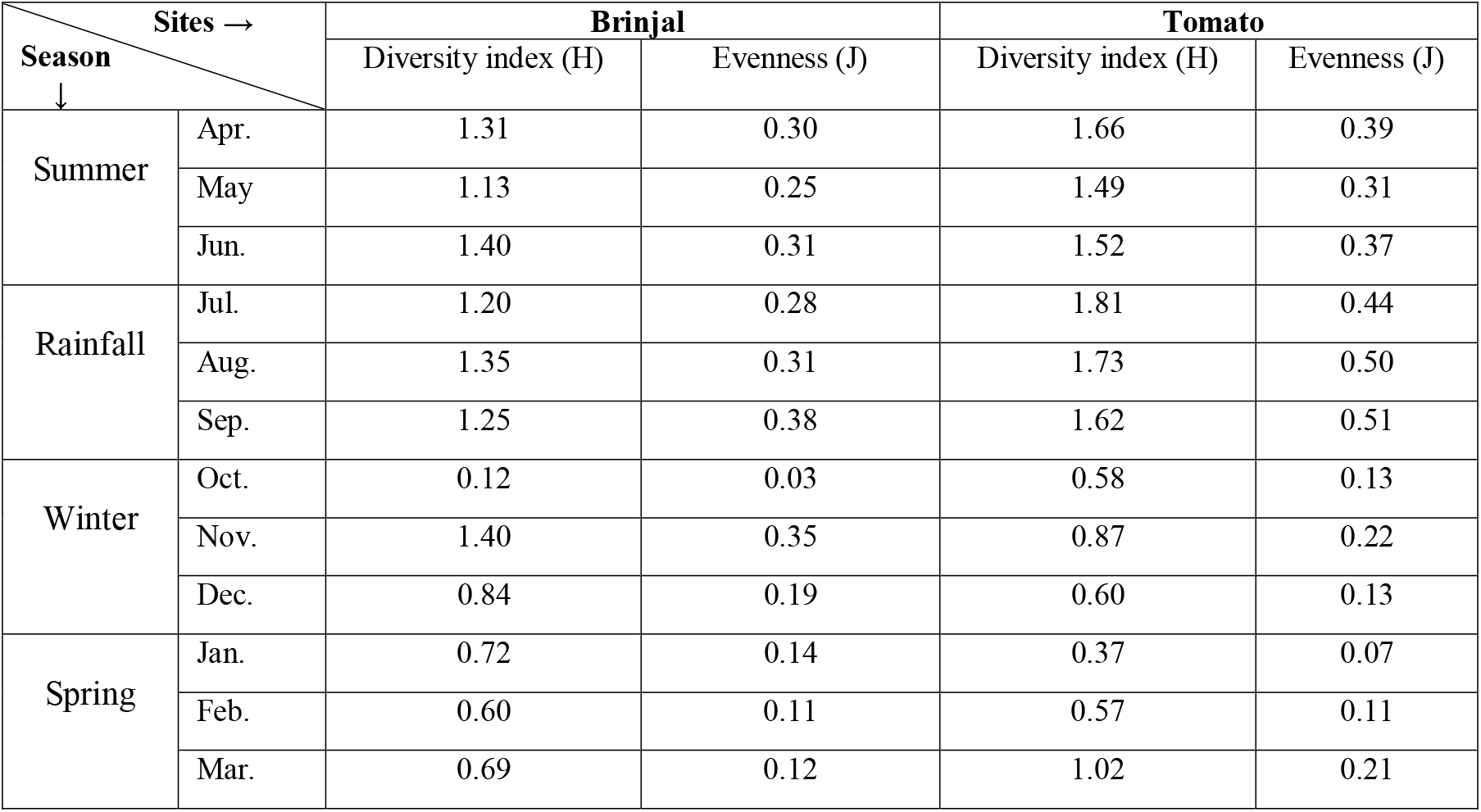
Seasonal diversity of soil microarthropods represented by Shannon Wiener diversity index (H) and Evenness (J) between two vegetational plots.

| Season<br>↓<br>Sites → |  | Brinjal |  | Tomato |  |
| --- | --- | --- | --- | --- | --- |
|  |  | Diversity index (H) | Evenness (J) | Diversity index (H) | Evenness (J) |
| Summer | Apr. | 1.31 | 0.30 | 1.66 | 0.39 |
|  | May | 1.13 | 0.25 | 1.49 | 0.31 |
|  | Jun. | 1.40 | 0.31 | 1.52 | 0.37 |
| Rainfall | Jul. | 1.20 | 0.28 | 1.81 | 0.44 |
|  | Aug. | 1.35 | 0.31 | 1.73 | 0.50 |
|  | Sep. | 1.25 | 0.38 | 1.62 | 0.51 |
| Winter | Oct. | 0.12 | 0.03 | 0.58 | 0.13 |
|  | Nov. | 1.40 | 0.35 | 0.87 | 0.22 |
|  | Dec. | 0.84 | 0.19 | 0.60 | 0.13 |
| Spring | Jan. | 0.72 | 0.14 | 0.37 | 0.07 |
|  | Feb. | 0.60 | 0.11 | 0.57 | 0.11 |
|  | Mar. | 0.69 | 0.12 | 1.02 | 0.21 |

The highly significant negative correlation was recorded between Collembolans population and soil temperature (r = −0.867, P<0.05), (figure 2) whereas available nitrogen was positively correlated (r = 0.847, P>0.05) in tomato plot. Both, Collembolans and Acari were most frequent (more than 80%) in brinjal plot, while soil dwelling Pterygotes (87.50%) in tomato plot (table 2).

**Figure: 2.**
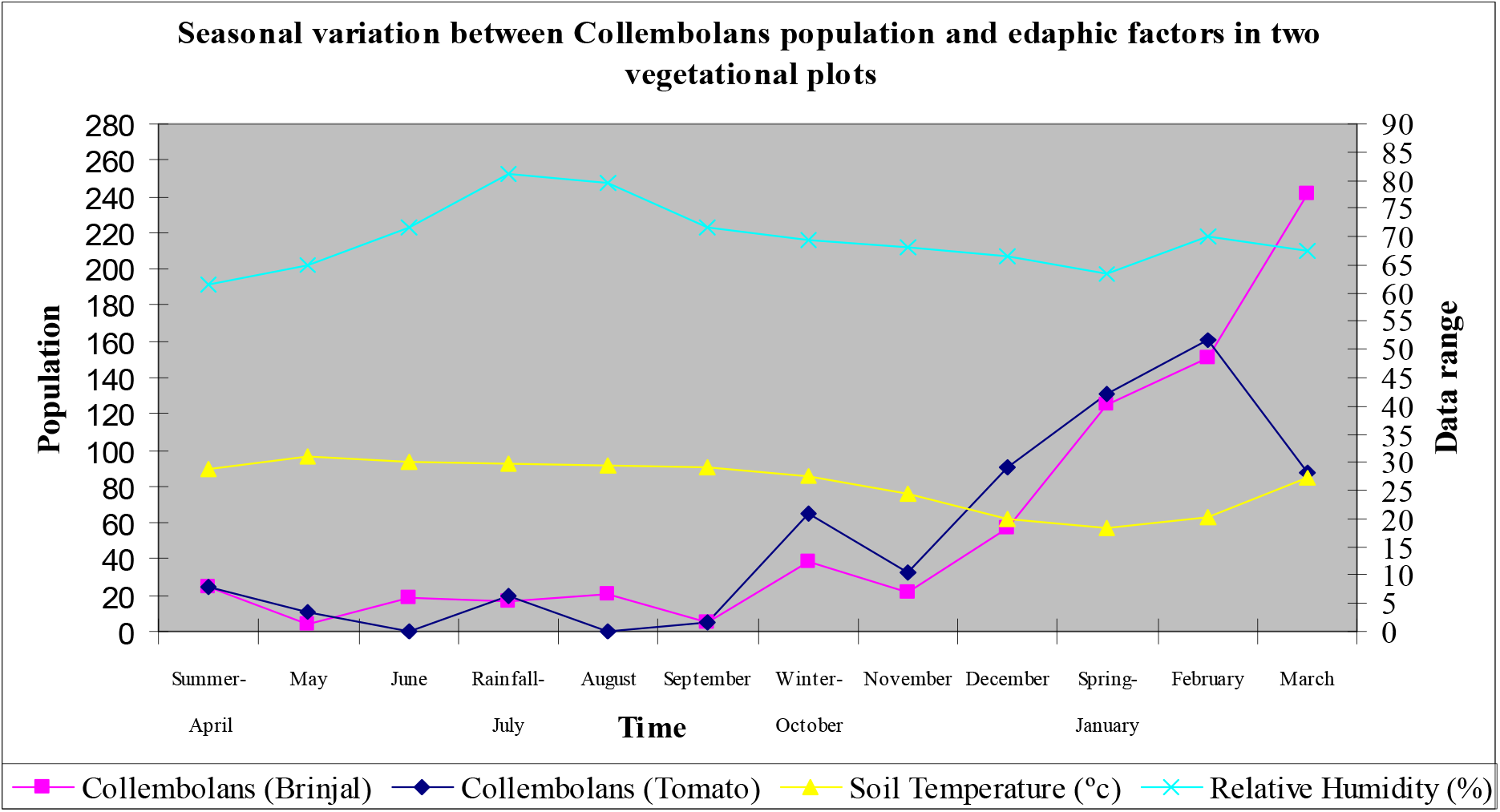
Seasonal variation between Collembolans population and edaphic factors in two vegetational plots at Aligarh.

## Discussion

Most of the earth’s arable land is already involved in some form of crop production to sustain the world populations (Altieri, 1994), while some agricultural practices and production systems have been evaluated for their short-term impact on sustainable production, few have been evaluated for their long-term impacts (Paoletti et al. 1993). Olfert et al. (2002) suggested that, cropping systems must incorporate the relationships between farm practices and the ecosystem used to create an equilibrium where farm inputs enhance rather than replace natural processes. Thus, soil microarthropods community is an important natural resource, helps on the nature and fertility of soil. On the other hand, habitat quality or vegetation cover is also another factor plays an important role in making the difference between communities and population of soil microarthropods. Insights the study, there was a significant difference in the population of soil microarthropods between the plots investigated whereas the plots situated at one sequence and soil of both plots was approximately same in nature and fertility of its production. However, some micro changes in edaphic factors have been recorded during study time.

Edaphic factors such as soil temperature, relative humidity, soil moisture and organic carbon most likely to influence organisms which are known to be sensitive to the farm practices, even for a limited period. Van Straalen (1997) also stated previously that, species richness and biological success of specific communities are positively related to the diversity of niches and soil microenvironments. It was noted that differences in dispersal rates of soil faunal species are likely to be strongly correlated with the differences in their population’s response and with other factors like temperature, soil moisture and organic matter of soil.

We have shown, brinjal plot was more diversified than compare to the tomato plot and overall density of soil microarthropods as well as Collembolans was greater in brinjal plot than compare to tomato plot. This may be due to the significant amount of litter present in the respective plot that was less recorded in tomato site. The peak population buildup of Collembola and Acarina during spring and winter months, while the sharp decline in summer months in both vegetational plots. In this regard, the reason is not yet clear; however, it may be due to cool temperatures with that of medium humid environment in winter months, because we have shown that as soon as the temperature increased the population of soil microarthropods decreased (figure 2). Thus, it can be said that temperature ranges are significantly correlated with soil microarthropods population dynamics. Ferguson and Joly (2002) also stated that, springtails and mite populations were regulated intrinsically by competition for food and secondarily by temperature as a function of climate.

Microclimatic differences due to vegetation cover between the plots investigated may be the other possibilities for differences in the soil microarthropods populations. Pankhurst (1997) also reported that the differences in soil microenvironments tremendously effect the soil microarthropods population. The possibility of population increased in the winter months, may be due to the stabilization of soil moisture contents in the respective season because the stabilize condition of soil moisture positively affect the population of soil microarthropods. It is well known fact that the soil moisture is an important edaphic factor to control the population dynamics of soil mesofauna.

One missing factor in our study was the litter quality comparison between the plots examined. However, brinjal plot was rich in terms of quantity of litter than compare to tomato plot. This may be the important reason for brinjal plot that was more diversified than tomato plot. The increased temperature and relative humidity, both were negatively correlated to the Collembolans population. We agree with the observations of various other researchers, as they concluded, the factors like soil moisture, temperature, soil pH, and food resources are known to affect the density of springtails (Altieri 1999, Chagnon et al. 2000, Olejniczak 2000, and Hasegawa 2001). Also we agree with the statement of the Snider and Butcher (1972) that they concluded, as temperature increases, the survival times of Collembola at specific relative humidity declines. However, the habitat quality or vegetation cover is most probably the responsible factor that affects the population buildup of soil inhabiting microarthropods. As shown in our observation, several other researchers have also concluded that changes in vegetation have been shown to change the composition of Collembola (Fox 1967, Moore et al. 1984, Fratello et al. 1985, Wardle et al. 1993).

The significant differences were recorded in the population of soil microarthropods between the plots investigated whereas the plots situated at one sequence and the distance between these two vegetational plots was only 15 meter as stated earlier. However, both Collembolans and Acari(mites) showed a consistent but similar trend of population fluctuation. So, it can be stated that the population of soil microarthropods vary plot to plot with time as a function of climate and vegetation cover affects the population of soil microarthropods significantly. Interestingly, seasonal differences have also pronounced that affect the particular group of soil microarthropods specifically on Collembolans.

## Conclusions

The changes in vegetation cover have been shown to change the composition of soil microarthropods. Collembolans and Acari(mites) showed a consistent but similar trend of fluctuations with time as a function of climate. The factors like soil moisture, temperature, relative humidity, pH of soil, and food resources, all have the cumulative effect on the density of soil microarthropods. Therefore, seasonal patterns are responsible for the population buildup of soil microarthropods. The cool temperatures with medium humid environment in winter months are the best environmental conditions for the survival and rich diversity of Collembolans.

